## Supplementary Materials for "Cerebellar and Subcortical Contributions to Working Memory Manipulation"

### 1. fMRIprep preprocessing

#### *Anatomical data preprocessing*

A total of 2 T1-weighted (T1w) images were found within the input BIDS dataset. All of them were corrected for intensity non-uniformity (INU) with `N4BiasFieldCorrection` [@n4], distributed with ANTs 2.3.1 [@ants, RRID:SCR_004757]. The T1w-reference was then skull-stripped with a *Nipype* implementation of the `antsBrainExtraction.sh` workflow (from ANTs), using OASIS30ANTs as target template. Brain tissue segmentation of cerebrospinal fluid (CSF), white-matter (WM) and gray-matter (GM) was performed on the brain-extracted T1w using `fast` [FSL 6.0.3:b862cdd5, RRID:SCR_002823, @fsl_fast]. An anatomical T1w-reference map was computed after registration of 2 T1w images (after INU-correction) using `mri_robust_template` [FreeSurfer 7.1.1, @fs_template]. Volume-based spatial normalization to two standard spaces (MNI152NLin6Asym, MNI152NLin2009cAsym) was performed through nonlinear registration with `antsRegistration` (ANTs 2.3.1), using brain-extracted versions of both T1w reference and the T1w template. The following templates were were selected for spatial normalization and accessed with *TemplateFlow* [23.0.0, @templateflow]: *FSL's MNI ICBM 152 non-linear 6th Generation Asymmetric Average Brain Stereotaxic Registration Model* [@mni152nlin6asym, RRID:SCR_002823; TemplateFlow ID: MNI152NLin6Asym], *ICBM 152 Nonlinear Asymmetrical template version 2009c* [@mni152nlin2009casym, RRID:SCR_008796; TemplateFlow ID: MNI152NLin2009cAsym].

#### *Functional data preprocessing*

For each of the 16 BOLD runs found per subject (across all tasks and sessions), the following preprocessing was performed. First, a reference volume and its skull-stripped version were generated using a custom methodology of *fMRIPrep*. Head-motion parameters with respect to the BOLD reference (transformation matrices, and six corresponding rotation and translation parameters) are estimated before any spatiotemporal filtering using `mcflirt` [FSL 6.0.3:b862cdd5, @mcflirt]. BOLD runs were slice-time corrected to 0.459s (0.5 of slice acquisition range 0s-0.917s) using `3dTshift` from AFNI 20170202 [@afni, RRID:SCR_005927].

The BOLD time-series (including slice-timing correction when applied) were resampled onto their original, native space by applying the transforms to correct for head-motion. These resampled BOLD time-series will be referred to as *preprocessed BOLD in original space*, or just *preprocessed BOLD*. The BOLD reference was then co-registered to the T1w reference using `mri_coreg` (FreeSurfer) followed by `flirt` [FSL 6.0.3:b862cdd5, @flirt] with the boundary-based registration [@bbr] cost-function. Co-registration was configured with six degrees of freedom. Several confounding time-series were calculated based on the *preprocessed BOLD*: framewise displacement (FD), DVARS and three region-wise global signals. FD was computed using two formulations following Power (absolute sum of relative motions, @power_fd_dvars) and Jenkinson (relative root mean square displacement between affines, @mcflirt). FD and DVARS are calculated for each functional run, both using their implementations in *Nipype* [following the definitions by @power_fd_dvars]. The three global signals are extracted within the CSF, the WM, and the whole-brain masks. Additionally, a set of physiological regressors were extracted to allow for component-based noise correction [*CompCor*, @compcor]. Principal components are estimated after high-pass filtering the *preprocessed BOLD* time-series (using a discrete cosine filter with 128s cut-off) for the two *CompCor* variants: temporal (tCompCor) and anatomical (aCompCor). tCompCor components are then calculated from the top 2% variable voxels within the brain mask. For aCompCor, three probabilistic masks (CSF, WM and combined CSF+WM) are generated in anatomical space. The implementation differs from that of Behzadi et al. in that instead of eroding the masks by 2 pixels on BOLD space, a mask of pixels that likely contain a volume fraction of GM is subtracted from the aCompCor masks. This mask is obtained by thresholding the corresponding partial volume map at 0.05, and it ensures components are not extracted from voxels containing a minimal fraction of GM. Finally, these masks are resampled into BOLD space and binarized by thresholding at 0.99 (as in the original implementation). Components are also calculated separately within the WM and CSF masks. For each CompCor decomposition, the *k* components with the largest singular values are retained, such that the retained components' time series are sufficient to explain 50 percent of variance across the nuisance mask (CSF, WM, combined, or temporal). The remaining components are dropped from consideration. The head-motion estimates calculated in the correction step were also placed within the corresponding confounds file. The confound time series derived from head motion estimates and global signals were expanded with the inclusion of temporal derivatives and quadratic terms for each [@confounds_satterthwaite_2013]. Frames that exceeded a threshold of 0.5 mm FD or 1.5

standardized DVARS were annotated as motion outliers. Additional nuisance timeseries are calculated by means of principal components analysis of the signal found within a thin band (*crown*) of voxels around the edge of the brain, as proposed by [@patriat_improved_2017].

The BOLD time-series were resampled into standard space, generating a *preprocessed BOLD run in MNI152NLin6Asym space*. First, a reference volume and its skull-stripped version were generated  using a custom methodology of *fMRIPrep*. All resamplings can be performed with *a single interpolation step* by composing all the pertinent transformations (i.e. head-motion transform matrices, susceptibility distortion correction when available, and co-registrations to anatomical and output spaces). Gridded (volumetric) resamplings were performed using `antsApplyTransforms` (ANTs), configured with Lanczos interpolation to minimize the smoothing effects of other kernels [@lanczos]. Non-gridded (surface) resamplings were performed using `mri_vol2surf (FreeSurfer).

Many internal operations of *fMRIPrep* use *Nilearn* 0.10.0 [@nilearn, RRID:SCR_001362], mostly within the functional processing workflow. For more details of the pipeline, see [the section corresponding to workflows in *fMRIPrep*'s documentation](https://fmriprep.readthedocs.io/en/latest/workflows.html "FMRIPrep's documentation").

### 2. LDA training and performance evaluation

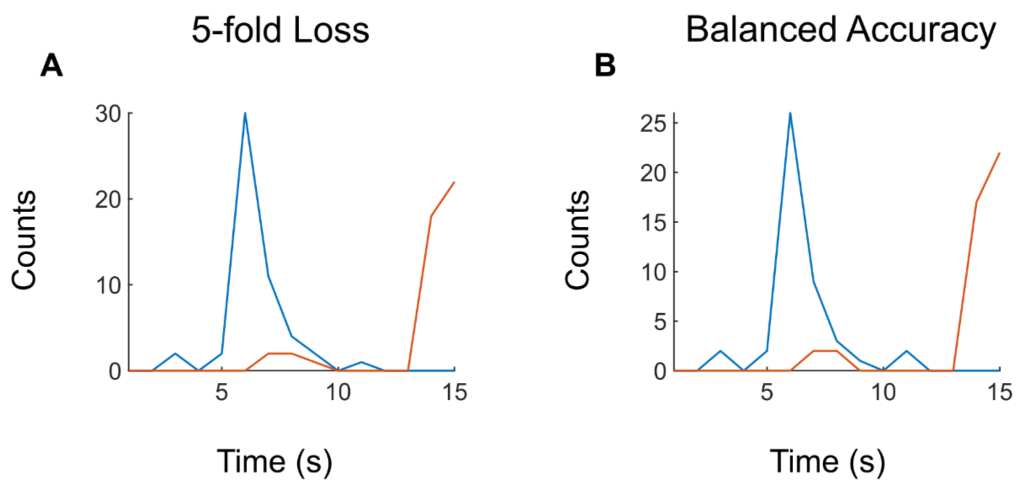

**Fig. S1.**

**Performance measures of LDA classifiers across time and with varying number of principal components. (A)** Number of times that each timepoint had the least loss. **(B)** Number of times that each timepoint had the highest balanced accuracy.

#### 3. Replication of LDA results in Schaefer 1000

Similar to the LDA training with the Schaefer 400 atlas, the timepoints which best separated the conditions were t = 6 (Easy vs. Hard), and t = 15 (Hard-Correct vs. Hard-Incorrect) (Figure 6). For these two separations, the minimum number of principal components (PC) in order to reach maximal balanced accuracy was 5 and 12 respectively. Following dimensionality reduction guidelines and choosing up to the first PC with added explained variance <1% (PC 12; Nguyen and Holmes, 2019). The first 12 PCs explained ~ 36% variance of the data.

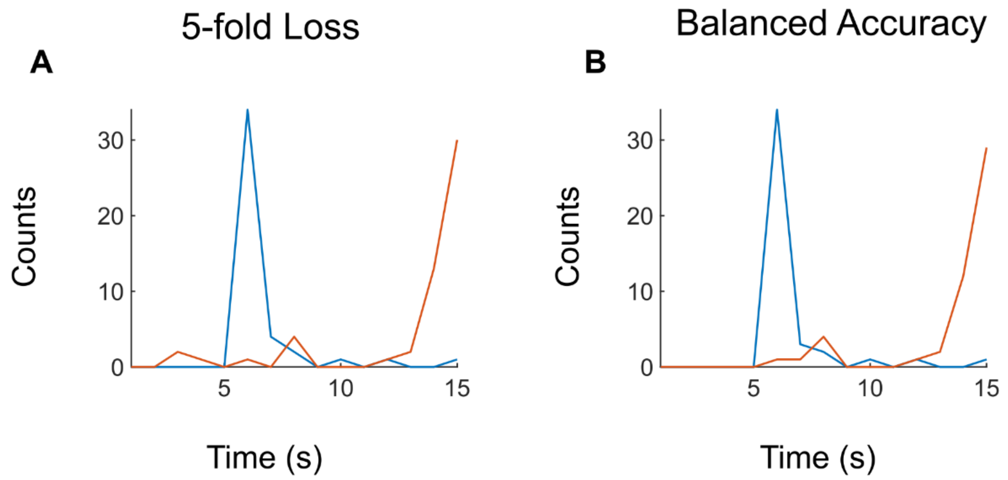

**Fig. S2.**

**Performance measures of LDA classifiers across time and with varying number of principal components for finer parcellation. (A)** Number of times that each timepoint had the least loss. **(B)** Number of times that each timepoint had the highest balanced accuracy.

### 4. Table 1. Labels for brain regions comparing Easy-Correct vs. Hard-Correct

| ROI Name | X | Y | Z |
|---|---|---|---|
| 17Networks_LH_VisCent_ExStr_1 | -36 | -62 | -17 |
| 17Networks_LH_VisCent_ExStr_2 | -23 | -73 | -10 |
| 17Networks_LH_VisCent_ExStr_6 | -41 | -87 | -3 |
| 17Networks_LH_VisCent_ExStr_7 | -46 | -73 | 6 |
| 17Networks_LH_VisCent_ExStr_9 | -39 | -84 | 14 |
| 17Networks_LH_VisCent_ExStr_10 | -11 | -97 | 17 |
| 17Networks_LH_VisCent_ExStr_11 | -25 | -85 | 21 |
| 17Networks_LH_VisPeri_ExStrInf_1 | -24 | -55 | -8 |
| 17Networks_LH_VisPeri_ExStrInf_2 | -12 | -62 | -5 |
| 17Networks_LH_VisPeri_ExStrInf_3 | -7 | -76 | -6 |
| 17Networks_LH_VisPeri_ExStrInf_4 | -13 | -43 | -5 |
| 17Networks_LH_VisPeri_ExStrInf_5 | -14 | -57 | 1 |
| 17Networks_LH_VisPeri_StriCal_1 | -5 | -88 | 2 |
| 17Networks_LH_VisPeri_StriCal_2 | -7 | -74 | 9 |
| 17Networks_LH_VisPeri_ExStrSup_1 | -19 | -65 | 7 |
| 17Networks_LH_VisPeri_ExStrSup_2 | -3 | -84 | 24 |
| 17Networks_LH_VisPeri_ExStrSup_3 | -12 | -71 | 20 |
| 17Networks_LH_VisPeri_ExStrSup_4 | -16 | -89 | 33 |
| 17Networks_LH_VisPeri_ExStrSup_5 | -12 | -81 | 36 |
| 17Networks_LH_SomMotA_1 | -8 | -15 | 47 |
| 17Networks_LH_SomMotA_4 | -48 | -29 | 58 |
| 17Networks_LH_SomMotA_5 | -39 | -25 | 53 |
| 17Networks_LH_SomMotA_6 | -9 | -38 | 54 |
| 17Networks_LH_SomMotA_7 | -4 | -25 | 56 |
| 17Networks_LH_SomMotA_8 | -4 | -9 | 59 |
| 17Networks_LH_SomMotA_11 | -30 | -38 | 65 |
| 17Networks_LH_SomMotA_12 | -23 | -11 | 65 |
| 17Networks_LH_SomMotA_19 | -12 | -27 | 73 |
| 17Networks_LH_SomMotB_Ins_1 | -36 | -24 | 10 |
| 17Networks_LH_SomMotB_S2_2 | -36 | -26 | 19 |
| 17Networks_LH_DorsAttnA_TempOcc_4 | -45 | -70 | -8 |
| 17Networks_LH_DorsAttnA_ParOcc_1 | -48 | -65 | 15 |
| 17Networks_LH_DorsAttnA_ParOcc_2 | -32 | -84 | 27 |
| 17Networks_LH_DorsAttnA_SPL_1 | -26 | -70 | 31 |
| 17Networks_LH_DorsAttnA_SPL_2 | -21 | -79 | 45 |
| 17Networks_LH_DorsAttnA_SPL_3 | -23 | -65 | 46 |
| 17Networks_LH_DorsAttnA_SPL_4 | -29 | -58 | 50 |
| 17Networks_LH_DorsAttnA_SPL_5 | -36 | -52 | 56 |
| 17Networks_LH_DorsAttnA_SPL_6 | -15 | -71 | 57 |

| | | | |
|---|---|---|---|
| 17Networks_LH_DorsAttnA_SPL_7 | -29 | -61 | 62 |
| 17Networks_LH_DorsAttnB_PostC_2 | -55 | -20 | 41 |
| 17Networks_LH_DorsAttnB_PostC_3 | -55 | -32 | 45 |
| 17Networks_LH_DorsAttnB_PostC_4 | -46 | -29 | 44 |
| 17Networks_LH_DorsAttnB_PostC_5 | -39 | -37 | 49 |
| 17Networks_LH_DorsAttnB_PostC_6 | -30 | -46 | 63 |
| 17Networks_LH_DorsAttnB_PostC_8 | -20 | -57 | 66 |
| 17Networks_LH_DorsAttnB_FEF_2 | -25 | -1 | 55 |
| 17Networks_LH_DorsAttnB_FEF_3 | -30 | -8 | 52 |
| 17Networks_LH_DorsAttnB_PrCv_1 | -50 | 3 | 38 |
| 17Networks_LH_SalVentAttnA_Ins_3 | -33 | 19 | 8 |
| 17Networks_LH_SalVentAttnA_FrMed_2 | -5 | 9 | 48 |
| 17Networks_LH_SalVentAttnB_PFCl_1 | -38 | 49 | 11 |
| 17Networks_LH_SalVentAttnB_Ins_1 | -34 | 16 | -8 |
| 17Networks_LH_SalVentAttnB_Ins_2 | -33 | 25 | -1 |
| 17Networks_LH_LimbicB_OFC_3 | -10 | 47 | -21 |
| 17Networks_LH_LimbicB_OFC_4 | -4 | 23 | -19 |
| 17Networks_LH_LimbicB_OFC_5 | -15 | 65 | -8 |
| 17Networks_LH_ContA_IPS_4 | -45 | -41 | 47 |
| 17Networks_LH_ContA_IPS_5 | -33 | -46 | 41 |
| 17Networks_LH_ContA_PFCd_1 | -21 | 5 | 65 |
| 17Networks_LH_ContA_PFCl_1 | -49 | 6 | 26 |
| 17Networks_LH_ContA_Cingm_1 | -3 | 5 | 29 |
| 17Networks_LH_ContB_IPL_1 | -49 | -60 | 47 |
| 17Networks_LH_ContB_IPL_3 | -42 | -52 | 49 |
| 17Networks_LH_ContB_PFCd_1 | -30 | 14 | 57 |
| 17Networks_LH_ContB_PFClv_1 | -42 | 49 | -6 |
| 17Networks_LH_ContB_PFCmp_1 | -4 | 28 | 47 |
| 17Networks_LH_ContC_pCun_2 | -9 | -77 | 45 |
| 17Networks_LH_DefaultA_IPL_1 | -47 | -64 | 31 |
| 17Networks_LH_DefaultA_PFCd_1 | -25 | 28 | 43 |
| 17Networks_LH_DefaultA_PFCd_2 | -18 | 36 | 48 |
| 17Networks_LH_DefaultA_PFCd_3 | -22 | 20 | 52 |
| 17Networks_LH_DefaultA_pCunPCC_1 | -4 | -53 | 20 |
| 17Networks_LH_DefaultA_pCunPCC_2 | -5 | -60 | 30 |
| 17Networks_LH_DefaultA_pCunPCC_3 | -7 | -44 | 32 |
| 17Networks_LH_DefaultA_PFCm_1 | -5 | 55 | -10 |
| 17Networks_LH_DefaultA_PFCm_2 | -6 | 35 | -9 |
| 17Networks_LH_DefaultA_PFCm_3 | -6 | 59 | 7 |
| 17Networks_LH_DefaultA_PFCm_4 | -6 | 45 | 6 |
| 17Networks_LH_DefaultA_PFCm_6 | -5 | 34 | 21 |
| 17Networks_LH_DefaultB_Temp_1 | -44 | 13 | -34 |
| 17Networks_LH_DefaultB_Temp_2 | -54 | -2 | -30 |

| | | | |
|---|---|---|---|
| 17Networks_LH_DefaultB_Temp_4 | -57 | -9 | -14 |
| 17Networks_LH_DefaultB_Temp_6 | -52 | -22 | -6 |
| 17Networks_LH_DefaultB_IPL_1 | -45 | -58 | 21 |
| 17Networks_LH_DefaultB_PFCd_1 | -4 | 51 | 28 |
| 17Networks_LH_DefaultB_PFCd_2 | -14 | 58 | 31 |
| 17Networks_LH_DefaultB_PFCd_4 | -8 | 43 | 51 |
| 17Networks_LH_DefaultB_PFCd_5 | -13 | 24 | 61 |
| 17Networks_LH_DefaultB_PFCd_6 | -6 | 10 | 65 |
| 17Networks_LH_DefaultB_PFCl_1 | -41 | 19 | 48 |
| 17Networks_LH_DefaultB_PFCv_1 | -36 | 22 | -16 |
| 17Networks_LH_DefaultB_PFCv_3 | -46 | 32 | -10 |
| 17Networks_LH_DefaultB_PFCv_5 | -53 | 19 | 11 |
| 17Networks_LH_DefaultC_IPL_1 | -40 | -79 | 30 |
| 17Networks_LH_DefaultC_Rsp_1 | -13 | -49 | 4 |
| 17Networks_LH_DefaultC_Rsp_2 | -8 | -52 | 9 |
| 17Networks_LH_DefaultC_Rsp_3 | -13 | -61 | 19 |
| 17Networks_LH_DefaultC_PHC_3 | -18 | -37 | -12 |
| 17Networks_RH_VisCent_ExStr_1 | 36 | -53 | -17 |
| 17Networks_RH_VisCent_ExStr_2 | 37 | -73 | -16 |
| 17Networks_RH_VisCent_ExStr_3 | 23 | -74 | -11 |
| 17Networks_RH_VisCent_Striate_1 | 8 | -92 | -2 |
| 17Networks_RH_VisCent_ExStr_7 | 35 | -89 | 2 |
| 17Networks_RH_VisCent_ExStr_9 | 43 | -79 | 10 |
| 17Networks_RH_VisCent_ExStr_10 | 13 | -94 | 19 |
| 17Networks_RH_VisCent_ExStr_11 | 27 | -87 | 21 |
| 17Networks_RH_VisPeri_ExStrInf_1 | 26 | -52 | -9 |
| 17Networks_RH_VisPeri_ExStrInf_2 | 18 | -36 | -12 |
| 17Networks_RH_VisPeri_ExStrInf_3 | 9 | -72 | -5 |
| 17Networks_RH_VisPeri_ExStrInf_4 | 13 | -58 | -3 |
| 17Networks_RH_VisPeri_ExStrInf_5 | 18 | -45 | -3 |
| 17Networks_RH_VisPeri_StriCal_1 | 9 | -74 | 9 |
| 17Networks_RH_VisPeri_StriCal_2 | 22 | -59 | 6 |
| 17Networks_RH_VisPeri_ExStrSup_1 | 16 | -66 | 19 |
| 17Networks_RH_VisPeri_ExStrSup_2 | 5 | -80 | 24 |
| 17Networks_RH_VisPeri_ExStrSup_3 | 14 | -78 | 34 |
| 17Networks_RH_VisPeri_ExStrSup_4 | 16 | -87 | 36 |
| 17Networks_RH_SomMotA_1 | 54 | -17 | 40 |
| 17Networks_RH_SomMotA_11 | 4 | -25 | 58 |
| 17Networks_RH_SomMotB_Cent_1 | 61 | 6 | 30 |
| 17Networks_RH_DorsAttnA_TempOcc_1 | 34 | -37 | -23 |
| 17Networks_RH_DorsAttnA_TempOcc_3 | 50 | -64 | -9 |
| 17Networks_RH_DorsAttnA_ParOcc_1 | 48 | -66 | 4 |
| 17Networks_RH_DorsAttnA_ParOcc_3 | 36 | -79 | 24 |

| | | | |
|---|---|---|---|
| 17Networks_RH_DorsAttnA_SPL_1 | 29 | -78 | 37 |
| 17Networks_RH_DorsAttnA_SPL_2 | 32 | -66 | 35 |
| 17Networks_RH_DorsAttnA_SPL_3 | 19 | -79 | 50 |
| 17Networks_RH_DorsAttnA_SPL_4 | 31 | -64 | 53 |
| 17Networks_RH_DorsAttnA_SPL_5 | 21 | -69 | 53 |
| 17Networks_RH_DorsAttnA_SPL_6 | 34 | -50 | 54 |
| 17Networks_RH_DorsAttnA_SPL_7 | 27 | -58 | 61 |
| 17Networks_RH_DorsAttnA_SPL_8 | 14 | -64 | 65 |
| 17Networks_RH_DorsAttnB_TempOcc_1 | 59 | -55 | -2 |
| 17Networks_RH_DorsAttnB_PostC_1 | 61 | -14 | 30 |
| 17Networks_RH_DorsAttnB_PostC_2 | 57 | -23 | 44 |
| 17Networks_RH_DorsAttnB_PostC_3 | 44 | -37 | 50 |
| 17Networks_RH_DorsAttnB_PostC_4 | 45 | -28 | 42 |
| 17Networks_RH_DorsAttnB_PostC_5 | 35 | -36 | 51 |
| 17Networks_RH_DorsAttnB_PostC_6 | 7 | -54 | 59 |
| 17Networks_RH_DorsAttnB_FEF_2 | 27 | -3 | 52 |
| 17Networks_RH_DorsAttnB_FEF_3 | 25 | -3 | 64 |
| 17Networks_RH_SalVentAttnA_PrC_1 | 51 | 3 | 41 |
| 17Networks_RH_SalVentAttnA_FrOper_3 | 54 | 12 | 12 |
| 17Networks_RH_SalVentAttnA_FrMed_2 | 6 | 11 | 58 |
| 17Networks_RH_SalVentAttnB_Ins_1 | 34 | 21 | -8 |
| 17Networks_RH_SalVentAttnB_Ins_2 | 37 | 23 | 5 |
| 17Networks_RH_SalVentAttnB_PFCmp_1 | 8 | 35 | 25 |
| 17Networks_RH_LimbicB_OFC_1 | 13 | 24 | -21 |
| 17Networks_RH_LimbicB_OFC_2 | 23 | 22 | -21 |
| 17Networks_RH_LimbicB_OFC_3 | 8 | 47 | -23 |
| 17Networks_RH_LimbicB_OFC_4 | 20 | 43 | -18 |
| 17Networks_RH_LimbicB_OFC_5 | 5 | 22 | -21 |
| 17Networks_RH_LimbicB_OFC_6 | 9 | 63 | -14 |
| 17Networks_RH_ContA_IPS_2 | 54 | -33 | 51 |
| 17Networks_RH_ContA_IPS_3 | 47 | -44 | 46 |
| 17Networks_RH_ContA_IPS_4 | 36 | -44 | 45 |
| 17Networks_RH_ContA_PFCl_1 | 50 | 30 | 18 |
| 17Networks_RH_ContA_PFCl_2 | 48 | 18 | 23 |
| 17Networks_RH_ContA_PFCl_3 | 47 | 29 | 28 |
| 17Networks_RH_ContA_PFCl_4 | 49 | 8 | 25 |
| 17Networks_RH_ContA_PFCl_5 | 39 | 11 | 34 |
| 17Networks_RH_ContA_Cingm_1 | 5 | 1 | 30 |
| 17Networks_RH_ContB_PFClv_3 | 42 | 51 | -6 |
| 17Networks_RH_ContB_PFCmp_1 | 5 | 28 | 48 |
| 17Networks_RH_DefaultA_pCunPCC_1 | 6 | -52 | 23 |
| 17Networks_RH_DefaultA_PFCm_1 | 5 | 41 | -11 |
| 17Networks_RH_DefaultA_PFCm_2 | 9 | 67 | 1 |

| | | | |
|---|---|---|---|
| 17Networks_RH_DefaultA_PFCm_3 | 7 | 42 | 4 |
| 17Networks_RH_DefaultA_PFCm_4 | 7 | 54 | 13 |
| 17Networks_RH_DefaultA_PFCm_6 | 6 | 25 | 18 |
| 17Networks_RH_DefaultB_PFCd_3 | 5 | 44 | 40 |
| 17Networks_RH_DefaultB_PFCd_4 | 14 | 39 | 52 |
| 17Networks_RH_DefaultB_PFCv_1 | 35 | 23 | -18 |
| 17Networks_RH_DefaultC_IPL_1 | 48 | -64 | 22 |
| 17Networks_RH_DefaultC_IPL_2 | 45 | -75 | 31 |
| 17Networks_RH_DefaultC_Rsp_1 | 14 | -46 | 4 |
| 17Networks_RH_DefaultC_Rsp_2 | 12 | -55 | 15 |
| 17Networks_RH_DefaultC_PHC_2 | 31 | -31 | -18 |
| HIP-head-m1-rh | 20 | -12 | 22 |
| THA-VAip-rh | 8 | -14 | 4 |
| THA-VAia-rh | 6 | -6 | 2 |
| HIP-head-l-rh | 30 | -14 | -20 |
| HIP-body-rh | 30 | -28 | -10 |
| THA-VPm-rh | 10 | -24 | 2 |
| PUT-VA-rh | 22 | 12 | -6 |
| PUT-DA-rh | 26 | 6 | 2 |
| PUT-VP-rh | 30 | -12 | 0 |
| PUT-DP-rh | 28 | -2 | 6 |
| CAU-VA-rh | 10 | 12 | 4 |
| CAU-DA-rh | 16 | 18 | 8 |
| CAU-body-rh | 14 | 6 | 16 |
| CAU-tail-rh | 16 | -2 | 20 |
| NAc-shell-rh | 12 | 10 | -6 |
| NAc-core-rh | 14 | 18 | -2 |
| pGP-rh | 24 | -8 | -2 |
| aGP-rh | 18 | 0 | -2 |
| HIP-head-m1-lh | -18 | -12 | -22 |
| THA-VAip-lh | -6 | -14 | 4 |
| THA-VAia-lh | -4 | -6 | 2 |
| HIP-body-lh | -28 | -28 | -10 |
| THA-VPm-lh | -8 | -24 | 2 |
| THA-VPl-lh | -16 | -22 | 4 |
| THA-DAl-lh | -14 | -20 | 12 |
| PUT-VA-lh | -20 | 12 | -6 |
| PUT-DA-lh | -24 | 6 | 2 |
| PUT-VP-lh | -28 | -12 | 0 |
| PUT-DP-lh | -26 | -2 | 6 |
| CAU-VA-lh | -8 | 12 | 4 |
| CAU-DA-lh | -14 | 18 | 8 |
| CAU-body-lh | -12 | 6 | 16 |

| | | | |
|---|---|---|---|
| CAU-tail-lh | -14 | -2 | 20 |
| NAc-shell-lh | -10 | 10 | -6 |
| NAc-core-lh | -12 | 18 | -2 |
| pGP-lh | -22 | -8 | -2 |
| aGP-lh | -16 | 0 | -2 |
| Left_I_IV | -4 | -50 | -11 |
| Right_I_IV | 4 | -50 | -11 |
| Right_V | 18 | -50 | -19 |
| Left_VI | -30 | -50 | -27 |
| Vermis_VI | 0 | -65 | -27 |
| Right_VI | 30 | -50 | -27 |
| Vermis_CrusII | 0 | -76 | -35 |
| Vermis_VIIb | 0 | -68 | -32 |
| Left_IX | -5 | -51 | -55 |
| Vermis_IX | 0 | -55 | -35 |
| Left_X | -22 | -35 | -43 |
| Vermis_X | 0 | -47 | -37 |

5. **Table 2. Labels for brain regions comparing hard-correct vs. hard-incorrect**

| ROI Name | X | Y | Z |
|---|---|---|---|
| 17Networks_LH_VisPeri_ExStrInf_2 | -12 | -62 | -5 |
| 17Networks_LH_VisPeri_ExStrInf_4 | -13 | -43 | -5 |
| 17Networks_LH_VisPeri_ExStrInf_5 | -14 | -57 | 1 |
| 17Networks_LH_VisPeri_StriCal_2 | -7 | -74 | 9 |
| 17Networks_LH_VisPeri_ExStrSup_1 | -19 | -65 | 7 |
| 17Networks_LH_VisPeri_ExStrSup_2 | -3 | -84 | 24 |
| 17Networks_LH_VisPeri_ExStrSup_3 | -12 | -71 | 20 |
| 17Networks_LH_VisPeri_ExStrSup_5 | -12 | -81 | 36 |
| 17Networks_LH_SomMotA_8 | -4 | -9 | 59 |
| 17Networks_LH_SomMotA_12 | -23 | -11 | 65 |
| 17Networks_LH_SomMotA_16 | -14 | -11 | 73 |
| 17Networks_LH_SomMotB_Cent_4 | -51 | -7 | 43 |
| 17Networks_LH_DorsAttnA_TempOcc_1 | -45 | -42 | -21 |
| 17Networks_LH_DorsAttnA_TempOcc_2 | -33 | -42 | -21 |
| 17Networks_LH_DorsAttnA_TempOcc_3 | -49 | -56 | -15 |
| 17Networks_LH_DorsAttnA_ParOcc_1 | -48 | -65 | 15 |
| 17Networks_LH_DorsAttnA_ParOcc_2 | -32 | -84 | 27 |
| 17Networks_LH_DorsAttnA_SPL_1 | -26 | -70 | 31 |
| 17Networks_LH_DorsAttnA_SPL_2 | -21 | -79 | 45 |
| 17Networks_LH_DorsAttnA_SPL_3 | -23 | -65 | 46 |
| 17Networks_LH_DorsAttnA_SPL_4 | -29 | -58 | 50 |
| 17Networks_LH_DorsAttnA_SPL_5 | -36 | -52 | 56 |
| 17Networks_LH_DorsAttnA_SPL_6 | -15 | -71 | 57 |
| 17Networks_LH_DorsAttnA_SPL_7 | -29 | -61 | 62 |
| 17Networks_LH_DorsAttnB_PostC_1 | -61 | -23 | 33 |
| 17Networks_LH_DorsAttnB_PostC_3 | -55 | -32 | 45 |
| 17Networks_LH_DorsAttnB_PostC_5 | -39 | -37 | 49 |
| 17Networks_LH_DorsAttnB_PostC_6 | -30 | -46 | 63 |
| 17Networks_LH_DorsAttnB_PostC_7 | -7 | -59 | 63 |
| 17Networks_LH_DorsAttnB_PostC_8 | -20 | -57 | 66 |
| 17Networks_LH_DorsAttnB_PostC_9 | -13 | -50 | 72 |
| 17Networks_LH_DorsAttnB_FEF_1 | -40 | -3 | 51 |
| 17Networks_LH_DorsAttnB_FEF_2 | -25 | -1 | 55 |
| 17Networks_LH_DorsAttnB_FEF_3 | -30 | -8 | 52 |
| 17Networks_LH_DorsAttnB_PrCv_1 | -50 | 3 | 38 |
| 17Networks_LH_SalVentAttnA_ParOper_2 | -58 | -44 | 27 |
| 17Networks_LH_SalVentAttnA_ParOper_3 | -61 | -36 | 33 |
| 17Networks_LH_SalVentAttnA_Ins_2 | -40 | -15 | -2 |
| 17Networks_LH_SalVentAttnA_Ins_3 | -33 | 19 | 8 |

| | | | |
|---|---|---|---|
| 17Networks_LH_SalVentAttnA_FrOper_2 | -52 | 9 | 13 |
| 17Networks_LH_SalVentAttnA_ParMed_1 | -11 | -27 | 41 |
| 17Networks_LH_SalVentAttnA_ParMed_2 | -13 | -41 | 47 |
| 17Networks_LH_SalVentAttnA_ParMed_3 | -6 | -49 | 57 |
| 17Networks_LH_SalVentAttnA_FrMed_1 | -7 | 0 | 41 |
| 17Networks_LH_SalVentAttnA_FrMed_2 | -5 | 9 | 48 |
| 17Networks_LH_SalVentAttnA_FrMed_3 | -8 | -3 | 71 |
| 17Networks_LH_SalVentAttnB_PFCl_2 | -29 | 43 | 30 |
| 17Networks_LH_SalVentAttnB_PFCl_3 | -36 | 32 | 38 |
| 17Networks_LH_SalVentAttnB_Ins_1 | -34 | 16 | -8 |
| 17Networks_LH_SalVentAttnB_Ins_2 | -33 | 25 | -1 |
| 17Networks_LH_SalVentAttnB_Ins_3 | -43 | 12 | 2 |
| 17Networks_LH_SalVentAttnB_OFC_1 | -27 | 49 | -14 |
| 17Networks_LH_SalVentAttnB_PFCmp_1 | -6 | 22 | 31 |
| 17Networks_LH_LimbicB_OFC_2 | -24 | 23 | -20 |
| 17Networks_LH_LimbicB_OFC_3 | -10 | 47 | -21 |
| 17Networks_LH_LimbicA_TempPole_1 | -37 | -5 | -42 |
| 17Networks_LH_LimbicA_TempPole_3 | -26 | -9 | -33 |
| 17Networks_LH_LimbicA_TempPole_5 | -40 | -21 | -27 |
| 17Networks_LH_LimbicA_TempPole_6 | -32 | 12 | -29 |
| 17Networks_LH_ContA_Temp_1 | -55 | -62 | -1 |
| 17Networks_LH_ContA_IPS_1 | -29 | -74 | 42 |
| 17Networks_LH_ContA_IPS_2 | -58 | -42 | 45 |
| 17Networks_LH_ContA_IPS_3 | -35 | -62 | 48 |
| 17Networks_LH_ContA_IPS_4 | -45 | -41 | 47 |
| 17Networks_LH_ContA_IPS_5 | -33 | -46 | 41 |
| 17Networks_LH_ContA_PFCd_1 | -21 | 5 | 65 |
| 17Networks_LH_ContA_PFClv_1 | -48 | 35 | 10 |
| 17Networks_LH_ContA_PFClv_2 | -42 | 38 | 22 |
| 17Networks_LH_ContA_PFCl_1 | -49 | 6 | 26 |
| 17Networks_LH_ContA_PFCl_2 | -45 | 20 | 27 |
| 17Networks_LH_ContA_PFCl_3 | -39 | 7 | 34 |
| 17Networks_LH_ContA_Cingm_1 | -3 | 5 | 29 |
| 17Networks_LH_ContB_Temp_2 | -60 | -49 | -10 |
| 17Networks_LH_ContB_IPL_2 | -53 | -50 | 45 |
| 17Networks_LH_ContB_PFCd_1 | -30 | 14 | 57 |
| 17Networks_LH_ContB_PFClv_2 | -28 | 58 | -1 |
| 17Networks_LH_ContB_PFClv_3 | -28 | 57 | 13 |
| 17Networks_LH_ContB_PFCmp_1 | -4 | 28 | 47 |
| 17Networks_LH_ContC_pCun_1 | -10 | -70 | 32 |
| 17Networks_LH_ContC_pCun_2 | -9 | -77 | 45 |
| 17Networks_LH_ContC_pCun_3 | -5 | -64 | 52 |
| 17Networks_LH_ContC_Cingp_1 | -6 | -41 | 24 |

| | | | |
|---|---|---|---|
| 17Networks_LH_ContC_Cingp_2 | -4 | -22 | 29 |
| 17Networks_LH_DefaultA_PFCd_1 | -25 | 28 | 43 |
| 17Networks_LH_DefaultA_PFCd_3 | -22 | 20 | 52 |
| 17Networks_LH_DefaultA_pCunPCC_2 | -5 | -60 | 30 |
| 17Networks_LH_DefaultA_pCunPCC_4 | -4 | -34 | 38 |
| 17Networks_LH_DefaultA_pCunPCC_6 | -3 | -68 | 41 |
| 17Networks_LH_DefaultA_pCunPCC_7 | -7 | -51 | 43 |
| 17Networks_LH_DefaultA_PFCm_2 | -6 | 35 | -9 |
| 17Networks_LH_DefaultA_PFCm_4 | -6 | 45 | 6 |
| 17Networks_LH_DefaultA_PFCm_6 | -5 | 34 | 21 |
| 17Networks_LH_DefaultB_Temp_1 | -44 | 13 | -34 |
| 17Networks_LH_DefaultB_Temp_2 | -54 | -2 | -30 |
| 17Networks_LH_DefaultB_Temp_4 | -57 | -9 | -14 |
| 17Networks_LH_DefaultB_Temp_5 | -61 | -35 | -3 |
| 17Networks_LH_DefaultB_Temp_6 | -52 | -22 | -6 |
| 17Networks_LH_DefaultB_IPL_1 | -45 | -58 | 21 |
| 17Networks_LH_DefaultB_IPL_2 | -57 | -55 | 30 |
| 17Networks_LH_DefaultB_PFCd_3 | -22 | 51 | 31 |
| 17Networks_LH_DefaultB_PFCd_6 | -6 | 10 | 65 |
| 17Networks_LH_DefaultB_PFCl_1 | -41 | 19 | 48 |
| 17Networks_LH_DefaultB_PFCl_2 | -42 | 7 | 48 |
| 17Networks_LH_DefaultB_PFCv_1 | -36 | 22 | -16 |
| 17Networks_LH_DefaultB_PFCv_2 | -36 | 37 | -13 |
| 17Networks_LH_DefaultB_PFCv_3 | -46 | 32 | -10 |
| 17Networks_LH_DefaultB_PFCv_4 | -48 | 28 | 0 |
| 17Networks_LH_DefaultB_PFCv_5 | -53 | 19 | 11 |
| 17Networks_LH_DefaultC_IPL_1 | -40 | -79 | 30 |
| 17Networks_LH_DefaultC_Rsp_1 | -13 | -49 | 4 |
| 17Networks_LH_DefaultC_Rsp_2 | -8 | -52 | 9 |
| 17Networks_LH_DefaultC_Rsp_3 | -13 | -61 | 19 |
| 17Networks_LH_DefaultC_PHC_2 | -30 | -33 | -18 |
| 17Networks_LH_TempPar_1 | -53 | 6 | -11 |
| 17Networks_LH_TempPar_2 | -61 | -13 | -3 |
| 17Networks_LH_TempPar_3 | -62 | -32 | 5 |
| 17Networks_LH_TempPar_4 | -52 | -43 | 5 |
| 17Networks_LH_TempPar_5 | -57 | -54 | 10 |
| 17Networks_LH_TempPar_6 | -59 | -49 | 16 |
| 17Networks_RH_VisPeri_ExStrInf_4 | 13 | -58 | -3 |
| 17Networks_RH_VisPeri_ExStrInf_5 | 18 | -45 | -3 |
| 17Networks_RH_VisPeri_StriCal_1 | 9 | -74 | 9 |
| 17Networks_RH_VisPeri_StriCal_2 | 22 | -59 | 6 |
| 17Networks_RH_VisPeri_ExStrSup_1 | 16 | -66 | 19 |
| 17Networks_RH_VisPeri_ExStrSup_2 | 5 | -80 | 24 |

| | | | |
|---|---|---|---|
| 17Networks_RH_VisPeri_ExStrSup_3 | 14 | -78 | 34 |
| 17Networks_RH_SomMotA_1 | 54 | -17 | 40 |
| 17Networks_RH_SomMotA_5 | 7 | -10 | 51 |
| 17Networks_RH_SomMotA_6 | 43 | -21 | 54 |
| 17Networks_RH_SomMotA_7 | 37 | -20 | 64 |
| 17Networks_RH_SomMotA_8 | 32 | -34 | 63 |
| 17Networks_RH_SomMotA_9 | 31 | -41 | 64 |
| 17Networks_RH_SomMotA_12 | 29 | -11 | 65 |
| 17Networks_RH_SomMotA_17 | 17 | -6 | 69 |
| 17Networks_RH_SomMotB_Ins_1 | 39 | -19 | 5 |
| 17Networks_RH_SomMotB_S2_4 | 41 | -29 | 18 |
| 17Networks_RH_SomMotB_S2_7 | 49 | -21 | 19 |
| 17Networks_RH_DorsAttnA_ParOcc_2 | 54 | -56 | 12 |
| 17Networks_RH_DorsAttnA_SPL_3 | 19 | -79 | 50 |
| 17Networks_RH_DorsAttnA_SPL_5 | 21 | -69 | 53 |
| 17Networks_RH_DorsAttnA_SPL_7 | 27 | -58 | 61 |
| 17Networks_RH_DorsAttnA_SPL_8 | 14 | -64 | 65 |
| 17Networks_RH_DorsAttnB_PostC_4 | 45 | -28 | 42 |
| 17Networks_RH_DorsAttnB_PostC_5 | 35 | -36 | 51 |
| 17Networks_RH_DorsAttnB_PostC_6 | 7 | -54 | 59 |
| 17Networks_RH_DorsAttnB_PostC_7 | 24 | -50 | 68 |
| 17Networks_RH_DorsAttnB_PostC_8 | 16 | -47 | 74 |
| 17Networks_RH_DorsAttnB_FEF_2 | 27 | -3 | 52 |
| 17Networks_RH_DorsAttnB_FEF_3 | 25 | -3 | 64 |
| 17Networks_RH_SalVentAttnA_ParOper_2 | 60 | -22 | 22 |
| 17Networks_RH_SalVentAttnA_ParOper_3 | 63 | -26 | 38 |
| 17Networks_RH_SalVentAttnA_Ins_1 | 40 | 5 | -15 |
| 17Networks_RH_SalVentAttnA_Ins_3 | 40 | -10 | -4 |
| 17Networks_RH_SalVentAttnA_FrMed_1 | 7 | 2 | 43 |
| 17Networks_RH_SalVentAttnA_ParMed_1 | 11 | -17 | 41 |
| 17Networks_RH_SalVentAttnA_ParMed_2 | 12 | -34 | 43 |
| 17Networks_RH_SalVentAttnA_FrMed_2 | 6 | 11 | 58 |
| 17Networks_RH_SalVentAttnA_ParMed_3 | 10 | -43 | 53 |
| 17Networks_RH_SalVentAttnA_ParMed_4 | 11 | -32 | 50 |
| 17Networks_RH_SalVentAttnA_FrMed_3 | 7 | -2 | 67 |
| 17Networks_RH_SalVentAttnA_FrMed_4 | 16 | 7 | 69 |
| 17Networks_RH_SalVentAttnB_IPL_1 | 62 | -37 | 37 |
| 17Networks_RH_SalVentAttnB_Ins_1 | 34 | 21 | -8 |
| 17Networks_RH_SalVentAttnB_Ins_2 | 37 | 23 | 5 |
| 17Networks_RH_SalVentAttnB_PFCmp_2 | 7 | 19 | 35 |
| 17Networks_RH_LimbicB_OFC_2 | 23 | 22 | -21 |
| 17Networks_RH_LimbicA_TempPole_2 | 49 | -7 | -39 |
| 17Networks_RH_LimbicA_TempPole_3 | 37 | 17 | -38 |

| | | | |
|---|---|---|---|
| 17Networks_RH_LimbicA_TempPole_4 | 39 | -15 | -31 |
| 17Networks_RH_LimbicA_TempPole_5 | 29 | 12 | -30 |
| 17Networks_RH_ContA_PFCd_1 | 24 | 10 | 58 |
| 17Networks_RH_ContA_PFCl_5 | 39 | 11 | 34 |
| 17Networks_RH_ContA_Cingm_1 | 5 | 1 | 30 |
| 17Networks_RH_ContB_IPL_1 | 55 | -45 | 33 |
| 17Networks_RH_ContB_PFCld_1 | 39 | 33 | 38 |
| 17Networks_RH_ContB_PFCmp_1 | 5 | 28 | 48 |
| 17Networks_RH_ContC_pCun_1 | 17 | -63 | 28 |
| 17Networks_RH_ContC_pCun_2 | 13 | -71 | 39 |
| 17Networks_RH_ContC_pCun_3 | 5 | -64 | 44 |
| 17Networks_RH_ContC_pCun_4 | 7 | -50 | 45 |
| 17Networks_RH_ContC_pCun_5 | 8 | -71 | 53 |
| 17Networks_RH_ContC_Cingp_1 | 7 | -44 | 20 |
| 17Networks_RH_ContC_Cingp_2 | 6 | -26 | 28 |
| 17Networks_RH_DefaultB_AntTemp_1 | 49 | 9 | -33 |
| 17Networks_RH_DefaultB_PFCv_1 | 35 | 23 | -18 |
| 17Networks_RH_DefaultB_PFCv_2 | 48 | 32 | -8 |
| 17Networks_RH_DefaultB_PFCv_3 | 54 | 24 | 6 |
| 17Networks_RH_DefaultC_IPL_2 | 45 | -75 | 31 |
| 17Networks_RH_DefaultC_Rsp_1 | 14 | -46 | 4 |
| 17Networks_RH_DefaultC_Rsp_2 | 12 | -55 | 15 |
| 17Networks_RH_TempPar_1 | 47 | 16 | -20 |
| 17Networks_RH_TempPar_3 | 49 | -20 | -8 |
| 17Networks_RH_TempPar_5 | 50 | -33 | 2 |
| 17Networks_RH_TempPar_6 | 59 | -46 | 7 |
| 17Networks_RH_TempPar_7 | 51 | -41 | 13 |
| 17Networks_RH_TempPar_9 | 55 | -46 | 19 |
| 17Networks_RH_TempPar_10 | 62 | -40 | 22 |
| HIP-head-l-rh | 30 | -14 | -20 |
| PUT-VA-rh | 22 | 12 | -6 |
| PUT-DA-rh | 26 | 6 | 2 |
| CAU-VA-rh | 10 | 12 | 4 |
| CAU-DA-rh | 16 | 18 | 8 |
| CAU-body-rh | 14 | 6 | 16 |
| NAc-shell-rh | 12 | 10 | -6 |
| NAc-core-rh | 14 | 18 | -2 |
| HIP-head-m1-lh | -18 | -12 | -22 |
| HIP-head-m2-lh | -20 | -18 | -16 |
| HIP-head-l-lh | -28 | -14 | -20 |
| HIP-body-lh | -28 | -28 | -10 |
| HIP-tail-lh | -22 | -38 | -2 |
| THA-VAs-lh | -8 | -10 | 12 |

| | | | |
|---|---|---|---|
| THA-DAm-lh | -8 | -24 | 12 |
| PUT-VA-lh | -20 | 12 | -6 |
| PUT-DA-lh | -24 | 6 | 2 |
| CAU-VA-lh | -8 | 12 | 4 |
| CAU-DA-lh | -14 | 18 | 8 |
| CAU-body-lh | -12 | 6 | 16 |
| CAU-tail-lh | -14 | -2 | 20 |
| lAMY-lh | -26 | -2 | -22 |
| NAc-shell-lh | -10 | 10 | -6 |
| NAc-core-lh | -12 | 18 | -2 |
| Left_VI | -30 | -50 | -27 |
| Vermis_VI | 0 | -65 | -27 |
| Right_VI | 30 | -50 | -27 |
| Right_CrusI | -45 | -65 | -32 |
| Right_CrusII | 39 | -65 | -47 |
| Right_VIIb | 33 | -65 | -53 |
| Right_IX | 5 | -51 | -55 |
| Right_X | 22 | -35 | -43 |

### 6. Table 3. Labels for brain regions important for both comparisons

| ROI Name | X | Y | Z |
|---|---|---|---|
| 17Networks_LH_VisPeri_ExStrInf_2 | -12 | -62 | -5 |
| 17Networks_LH_VisPeri_ExStrInf_4 | -13 | -43 | -5 |
| 17Networks_LH_VisPeri_ExStrInf_5 | -14 | -57 | 1 |
| 17Networks_LH_VisPeri_StriCal_2 | -7 | -74 | 9 |
| 17Networks_LH_VisPeri_ExStrSup_1 | -19 | -65 | 7 |
| 17Networks_LH_VisPeri_ExStrSup_2 | -3 | -84 | 24 |
| 17Networks_LH_VisPeri_ExStrSup_3 | -12 | -71 | 20 |
| 17Networks_LH_VisPeri_ExStrSup_5 | -12 | -81 | 36 |
| 17Networks_LH_SomMotA_8 | -4 | -9 | 59 |
| 17Networks_LH_SomMotA_12 | -23 | -11 | 65 |
| 17Networks_LH_DorsAttnA_ParOcc_1 | -48 | -65 | 15 |
| 17Networks_LH_DorsAttnA_ParOcc_2 | -32 | -84 | 27 |
| 17Networks_LH_DorsAttnA_SPL_1 | -26 | -70 | 31 |
| 17Networks_LH_DorsAttnA_SPL_2 | -21 | -79 | 45 |
| 17Networks_LH_DorsAttnA_SPL_3 | -23 | -65 | 46 |
| 17Networks_LH_DorsAttnA_SPL_4 | -29 | -58 | 50 |
| 17Networks_LH_DorsAttnA_SPL_5 | -36 | -52 | 56 |
| 17Networks_LH_DorsAttnA_SPL_6 | -15 | -71 | 57 |
| 17Networks_LH_DorsAttnA_SPL_7 | -29 | -61 | 62 |
| 17Networks_LH_DorsAttnB_PostC_3 | -55 | -32 | 45 |
| 17Networks_LH_DorsAttnB_PostC_5 | -39 | -37 | 49 |
| 17Networks_LH_DorsAttnB_PostC_6 | -30 | -46 | 63 |
| 17Networks_LH_DorsAttnB_PostC_8 | -20 | -57 | 66 |
| 17Networks_LH_DorsAttnB_FEF_2 | -25 | -1 | 55 |
| 17Networks_LH_DorsAttnB_FEF_3 | -30 | -8 | 52 |
| 17Networks_LH_DorsAttnB_PrCv_1 | -50 | 3 | 38 |
| 17Networks_LH_SalVentAttnA_Ins_3 | -33 | 19 | 8 |
| 17Networks_LH_SalVentAttnA_FrMed_2 | -5 | 9 | 48 |
| 17Networks_LH_SalVentAttnB_Ins_1 | -34 | 16 | -8 |
| 17Networks_LH_SalVentAttnB_Ins_2 | -33 | 25 | -1 |
| 17Networks_LH_LimbicB_OFC_3 | -10 | 47 | -21 |
| 17Networks_LH_ContA_IPS_4 | -45 | -41 | 47 |
| 17Networks_LH_ContA_IPS_5 | -33 | -46 | 41 |
| 17Networks_LH_ContA_PFCd_1 | -21 | 5 | 65 |
| 17Networks_LH_ContA_PFCl_1 | -49 | 6 | 26 |
| 17Networks_LH_ContA_Cingm_1 | -3 | 5 | 29 |
| 17Networks_LH_ContB_PFCd_1 | -30 | 14 | 57 |
| 17Networks_LH_ContB_PFCmp_1 | -4 | 28 | 47 |
| 17Networks_LH_ContC_pCun_2 | -9 | -77 | 45 |
| 17Networks_LH_DefaultA_PFCd_1 | -25 | 28 | 43 |
| 17Networks_LH_DefaultA_PFCd_3 | -22 | 20 | 52 |

| | | | |
|---|---|---|---|
| 17Networks_LH_DefaultA_pCunPCC_2 | -5 | -60 | 30 |
| 17Networks_LH_DefaultA_PFCm_2 | -6 | 35 | -9 |
| 17Networks_LH_DefaultA_PFCm_4 | -6 | 45 | 6 |
| 17Networks_LH_DefaultA_PFCm_6 | -5 | 34 | 21 |
| 17Networks_LH_DefaultB_Temp_1 | -44 | 13 | -34 |
| 17Networks_LH_DefaultB_Temp_2 | -54 | -2 | -30 |
| 17Networks_LH_DefaultB_Temp_4 | -57 | -9 | -14 |
| 17Networks_LH_DefaultB_Temp_6 | -52 | -22 | -6 |
| 17Networks_LH_DefaultB_IPL_1 | -45 | -58 | 21 |
| 17Networks_LH_DefaultB_PFCd_6 | -6 | 10 | 65 |
| 17Networks_LH_DefaultB_PFCl_1 | -41 | 19 | 48 |
| 17Networks_LH_DefaultB_PFCv_1 | -36 | 22 | -16 |
| 17Networks_LH_DefaultB_PFCv_3 | -46 | 32 | -10 |
| 17Networks_LH_DefaultB_PFCv_5 | -53 | 19 | 11 |
| 17Networks_LH_DefaultC_IPL_1 | -40 | -79 | 30 |
| 17Networks_LH_DefaultC_Rsp_1 | -13 | -49 | 4 |
| 17Networks_LH_DefaultC_Rsp_2 | -8 | -52 | 9 |
| 17Networks_LH_DefaultC_Rsp_3 | -13 | -61 | 19 |
| 17Networks_RH_VisPeri_ExStrInf_4 | 13 | -58 | -3 |
| 17Networks_RH_VisPeri_ExStrInf_5 | 18 | -45 | -3 |
| 17Networks_RH_VisPeri_StriCal_1 | 9 | -74 | 9 |
| 17Networks_RH_VisPeri_StriCal_2 | 22 | -59 | 6 |
| 17Networks_RH_VisPeri_ExStrSup_1 | 16 | -66 | 19 |
| 17Networks_RH_VisPeri_ExStrSup_2 | 5 | -80 | 24 |
| 17Networks_RH_VisPeri_ExStrSup_3 | 14 | -78 | 34 |
| 17Networks_RH_SomMotA_1 | 54 | -17 | 40 |
| 17Networks_RH_DorsAttnA_SPL_3 | 19 | -79 | 50 |
| 17Networks_RH_DorsAttnA_SPL_5 | 21 | -69 | 53 |
| 17Networks_RH_DorsAttnA_SPL_7 | 27 | -58 | 61 |
| 17Networks_RH_DorsAttnA_SPL_8 | 14 | -64 | 65 |
| 17Networks_RH_DorsAttnB_PostC_4 | 45 | -28 | 42 |
| 17Networks_RH_DorsAttnB_PostC_5 | 35 | -36 | 51 |
| 17Networks_RH_DorsAttnB_PostC_6 | 7 | -54 | 59 |
| 17Networks_RH_DorsAttnB_FEF_2 | 27 | -3 | 52 |
| 17Networks_RH_DorsAttnB_FEF_3 | 25 | -3 | 64 |
| 17Networks_RH_SalVentAttnA_FrMed_2 | 6 | 11 | 58 |
| 17Networks_RH_SalVentAttnB_Ins_1 | 34 | 21 | -8 |
| 17Networks_RH_SalVentAttnB_Ins_2 | 37 | 23 | 5 |
| 17Networks_RH_LimbicB_OFC_2 | 23 | 22 | -21 |
| 17Networks_RH_ContA_PFCl_5 | 39 | 11 | 34 |
| 17Networks_RH_ContA_Cingm_1 | 5 | 1 | 30 |
| 17Networks_RH_ContB_PFCmp_1 | 5 | 28 | 48 |
| 17Networks_RH_DefaultB_PFCv_1 | 35 | 23 | -18 |

| | | | |
|---|---|---|---|
| 17Networks_RH_DefaultC_IPL_2 | 45 | -75 | 31 |
| 17Networks_RH_DefaultC_Rsp_1 | 14 | -46 | 4 |
| 17Networks_RH_DefaultC_Rsp_2 | 12 | -55 | 15 |
| HIP-head-l-rh | 30 | -14 | -20 |
| PUT-VA-rh | 22 | 12 | -6 |
| PUT-DA-rh | 26 | 6 | 2 |
| CAU-VA-rh | 10 | 12 | 4 |
| CAU-DA-rh | 16 | 18 | 8 |
| CAU-body-rh | 14 | 6 | 16 |
| NAc-shell-rh | 12 | 10 | -6 |
| NAc-core-rh | 14 | 18 | -2 |
| HIP-head-m1-lh | -18 | -12 | -22 |
| HIP-body-lh | -28 | -28 | -10 |
| PUT-VA-lh | -20 | 12 | -6 |
| PUT-DA-lh | -24 | 6 | 2 |
| CAU-VA-lh | -8 | 12 | 4 |
| CAU-DA-lh | -14 | 18 | 8 |
| CAU-body-lh | -12 | 6 | 16 |
| CAU-tail-lh | -14 | -2 | 20 |
| NAc-shell-lh | -10 | 10 | -6 |
| NAc-core-lh | -12 | 18 | -2 |
| Left_VI | -30 | -50 | -27 |
| Vermis_VI | 0 | -65 | -27 |
| Right_VI | 30 | -50 | -27 |

### 7. Brain maps for other quadrants

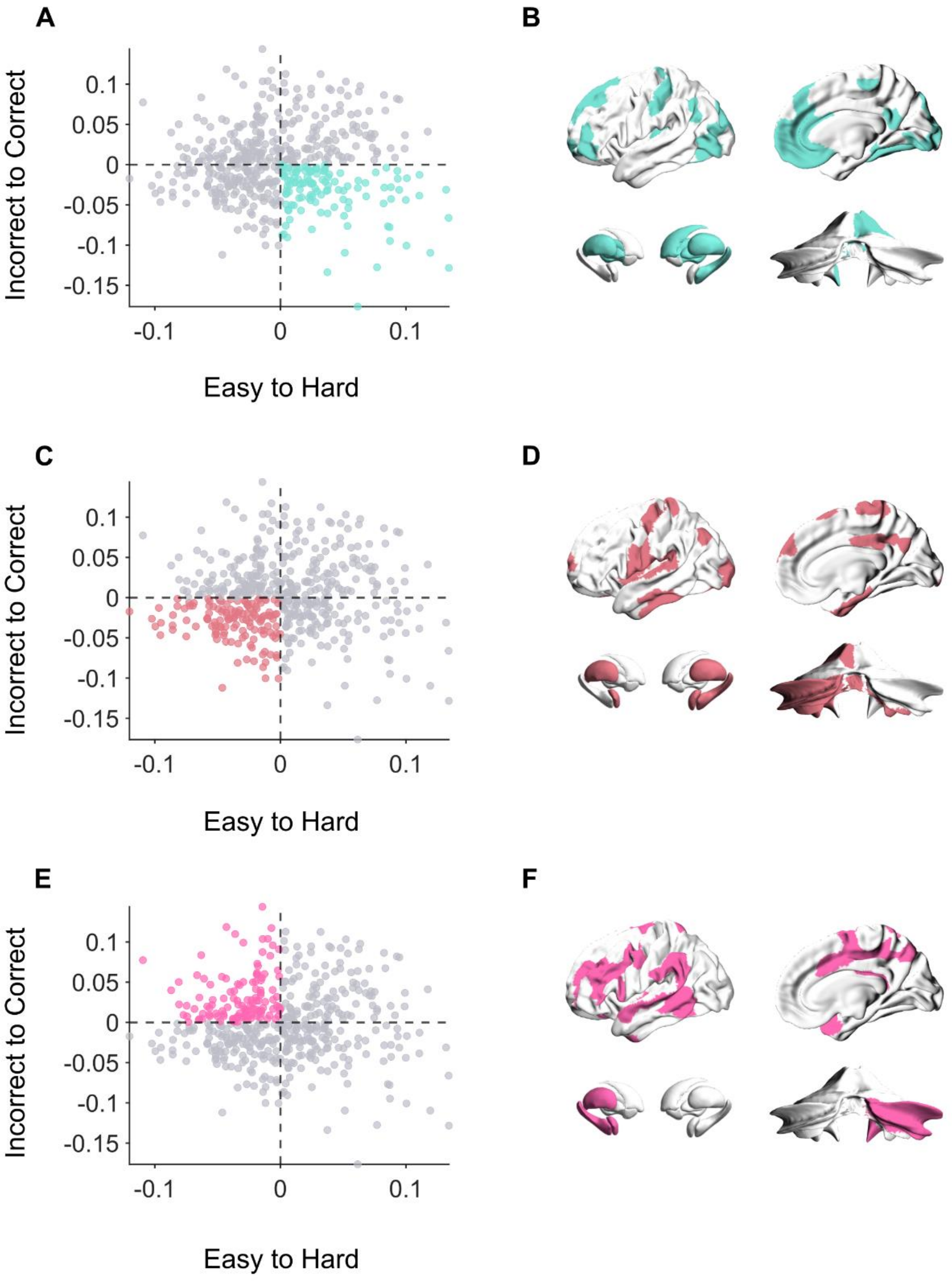

**Fig. S3.**

**Brain maps for other quadrants. (A, B)** Quadrant 2: regions related to Hard-Incorrect trials. **(C, D)** Quadrant 3: regions related to Easy-Incorrect trials. **(E, F)** Quadrant 4: regions related to Easy-Correct trials.
